# Novel autotrophic organisms contribute significantly to the internal carbon cycling potential of a boreal lake

**DOI:** 10.1101/309633

**Authors:** Sari Peura, Moritz Buck, Sanni L Aalto, Sergio E. Morales, Hannu Nykänen, Alexander Eiler

## Abstract

The authors declare no conflict of interest

Funding sources: the Academy of Finland, Science for Life Laboratories, Tryggers Foundation, the Swedish Research Council VR and the Swedish Foundation for strategic research

**Abstract:** Oxygen stratified lakes are typical for the boreal zone, and also a major source of greenhouse gas emissions in the region. Due to shallow light penetration, restricting the growth of phototrophic organisms, and large allochthonous organic carbon inputs from the catchment area, the lake metabolism is expected to be dominated by heterotrophic organisms. In this study we test this assumption and show that the potential for autotrophic carbon fixation and internal carbon cycling is high throughout the water column. Further, we show that during the summer stratification carbon fixation can exceed respiration in a boreal lake even below the euphotic zone. Metagenome assembled genomes and 16S profiling of a vertical transect of the lake revealed multiple organisms in oxygen depleted compartment belonging to novel or poorly characterized phyla. Many of these organisms were chemolithotrophic, deriving their energy from reactions related to sulfur, iron and nitrogen transformations. The community as well as the functions were stratified following the redox potentials. The autotrophic potential in the lake metagenome below the oxygenic zone was high, pointing towards a need for revising our concepts of internal carbon cycling in boreal lakes. Further, the importance of chemolithoautotrophy for the internal carbon cycling suggests that many predicted climate change associated changes in the physical properties of the lake, such as altered mixing patterns, likely have consequences for the whole lake metabolism even beyond the impact to the phototrophic community.

**Importance:** Autotrophic organisms at the base of the food web are the only life form capable of turning inorganic carbon into organic form, facilitating the survival of all other organisms. In certain environments the autotrophic production is limited by environmental conditions and the food web is supported by carbon coming from outside the ecosystem. One such environment is stratified boreal lakes, which are
one of the biggest sources of greenhouse gas emissions in the boreal region. Thus, carbon cycling in these habitats is of outmost importance for the future climate. Here we demonstrate a high potential for internal carbon cycling via phototrophic and novel chemolithotrophic organisms in the dark and anoxic layers of a boreal lake. Our results significantly increase our knowledge on the microbial communities and their metabolic potential in oxygen depleted freshwaters and help to understand and predict how climate change induced alterations could impact the lake carbon dynamics.

## Introduction

All life on Earth depends on carbon fixation, where microorganisms convert inorganic carbon dioxide into organic compounds and living biomass. Currently, oxygenic phototrophs, deriving their energy from the sun light, are regarded as the most important carbon fixers. However, on planetary time scales anoxygenic phototrophs and chemotrophs have been more prevalent (1). Even today chemolithoautotrophy is a major strategy in many environments, such as deep-sea vents and sediments (2-4). In these conditions carbon fixation is driven by a redox gradient (i.e. biogeochemical gradient of reductants and oxidants sorted by their redox potentials) often located in the border zone between oxic and anoxic conditions. These redox transition zones constitute a major share of the Earth’s biosphere, and have a significant impact on surrounding entities, such as elemental cycles and food webs (5-7).

Chemolithotrophy may have a large impact on the carbon cycle in many environments. For example, in boreal lakes there is a steep redox gradient at the oxic-anoxic border, which could facilitate carbon fixation via chemolithotrophy. Still, our knowledge on the chemolithotrophic energy generation in these habitats is poor. Some first studies have highlighted the importance of chemolithotrophy for carbon assimilation in freshwater lakes such as Kivu, a freshwater lake in central Africa with high methane (CH_4_) concentration (2). Furthermore, two recent studies have shown genetic potential for autotrophy in microbial communities in the anoxic water masses of boreal lakes (8, 9). This suggests that chemolithotrophy could be a common process fueling assimilation of inorganic carbon in boreal lakes. Thus, dark carbon fixation may play a significant role in the internal carbon cycling, and thus, modulate lake food webs and ultimately whole lake carbon balances.

Small boreal lakes are important drivers of global greenhouse gas (GHG) emissions (10, 11) and their importance was highlighted in the recent report of the IPCC (12). Globally, water bodies smaller than 0.001 km^2^ contribute 40% of all methane emissions from inland waters (13). Typically, these small lakes and ponds are characterized by high concentrations of dissolved organic carbon (DOC) and shallow light penetration depth, which leads to steep stratification of oxygen and other electron acceptors and donors through most of the year. The stratification coincides with a distinct set of bacterial and archaeal phyla organized according to the vertical redox gradient (9, 14, 15). The microbial communities in these lakes may harbor organisms that have the potential for photoautotrophy under low light intensity (14, 16, 17) and for chemoautotrophy throughout the water column (9). These predictions are based on taxonomic information of 16S rRNA genes combined with functional gene inventories and genomic data of related cultivated representatives, rather than a true reconstruction of the genetic makeup of individual organisms or whole communities along the redox tower. This lack of a detailed metabolic picture limits our understanding of the functional potential of microbes in boreal lakes.

We studied the potential of the microbial community for chemoautotrophy in lake Alinen Mustajärvi, a well-characterized boreal lake located in southern Finland (15) (18). This lake exhibits the typical characteristic features of boreal lakes including: 1) a high load of terrestrial organic carbon resulting in net heterotrophy of the system (18), 2) a gradient of oxygen, temperature and light, and 3) stratification of the microbial community (9, 14, 15). We combined dark carbon fixation measurements with a survey of the functional potential of the microbial community (shotgun sequencing of the total DNA) from a vertical transect of the lake water column. Our aim was to link lake chemistry to the prevalence of genes related to energy generation via redox reactions and inorganic carbon assimilation. Moreover, we used annotated metagenome-assembled genomes (MAGs) to obtain metabolic reconstructions of uncharacterized and novel lake microbes and to identify the key chemoautotrophs in the lake. Our hypotheses were that i) the chemical stratification of the lake was linked to microbial activity and would be reflected in the functional potential and structure of the communities, ii) the lake harbors an abundant and diverse chemoautotrophic community, and iii) chemoautotrophic pathways would be enriched below the euphotic zone of the lake, leading to a high potential for internal carbon cycling.

## Results and Discussion

### Microbial community structure and CO_2_ incorporation in the water column

The metabolic pathways and organisms that could be involved in autotrophic processes in the water column of lake Alinen Mustajärvi were studied based on metagenomic (shotgun sequencing and 16S rRNA gene amplicons) and geochemical analyses from samples taken in 2013. This vertical transect of the lake covered 13 depths. Additionally, the community composition was surveyed in 2008 using aclone library covering 4 depths and 303 clones of 16S rRNA genes. We also measured inorganic carbon dynamics biweekly in 2008, monthly in 2009 and on 3 occasions in 2010. The carbon measurements indicated that at 3 meters depth the incorporation of inorganic carbon dominated over respiration during parts of the ice-free season (Fig. 1, Fig. S1). Time of net autotrophy coincided with stratification of the lake and vertical structuring of the microbial community along the vertical gradient (Fig. 2A). The maximum value for dark carbon assimilation at 3 m depth was 10 % of the average net primary production (carbon assimilation in the light) within the euphotic zone (Fig. 1, (18)). While these are point measurements and should be taken with caution, they are well in line with dark carbon fixation values measured from lake sediments (19).

**Figure 1.**
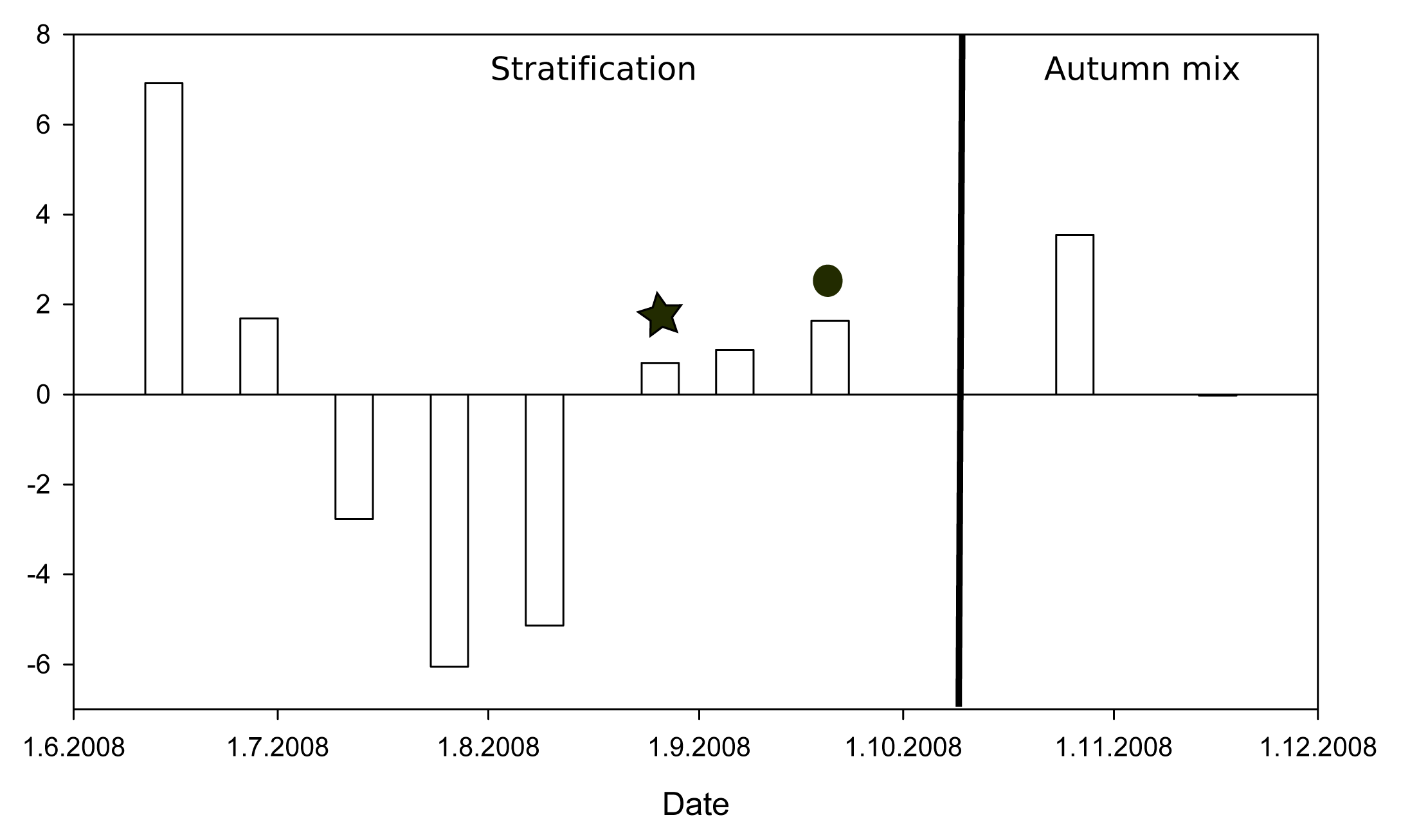
Change in CO_2_ concentration over 24 h *in situ* dark incubations during open water season 2008 at a depth of 3 m. Positive values indicate respiration dominating over CO_2_ assimilation while negative values show net incorporation of inorganic carbon into biomass. Star depicts the time point the clone library was retrieved in 2008 and the circle the time point when the amplicon and shot gun samples were taken in 2013.

The analysis of 16S rRNA genes indicated that the communities were dominated by similar phyla in 2008 and 2013 (Fig. 2A-B) and the community composition was consistent with previously published profiles from the same lake (15).*Actinobacteria* and *Alphaproteobacteria* were the major community members at the surface layer (epilimnion) and *Chlorobia* in the anoxic bottom layer (hypolimnion). In 2013 another major community member in the hypolimnion was the Parcubacteria, which was missing from the community in 2008 due to a mismatch in the primers used at the time (15). Community patterns were conserved across methods within the same year (Correlation in a symmetric Procrustes rotation of the 16S rRNA and metagenome communities in 2013 0.962, p>0.001).

**Figure 2.**
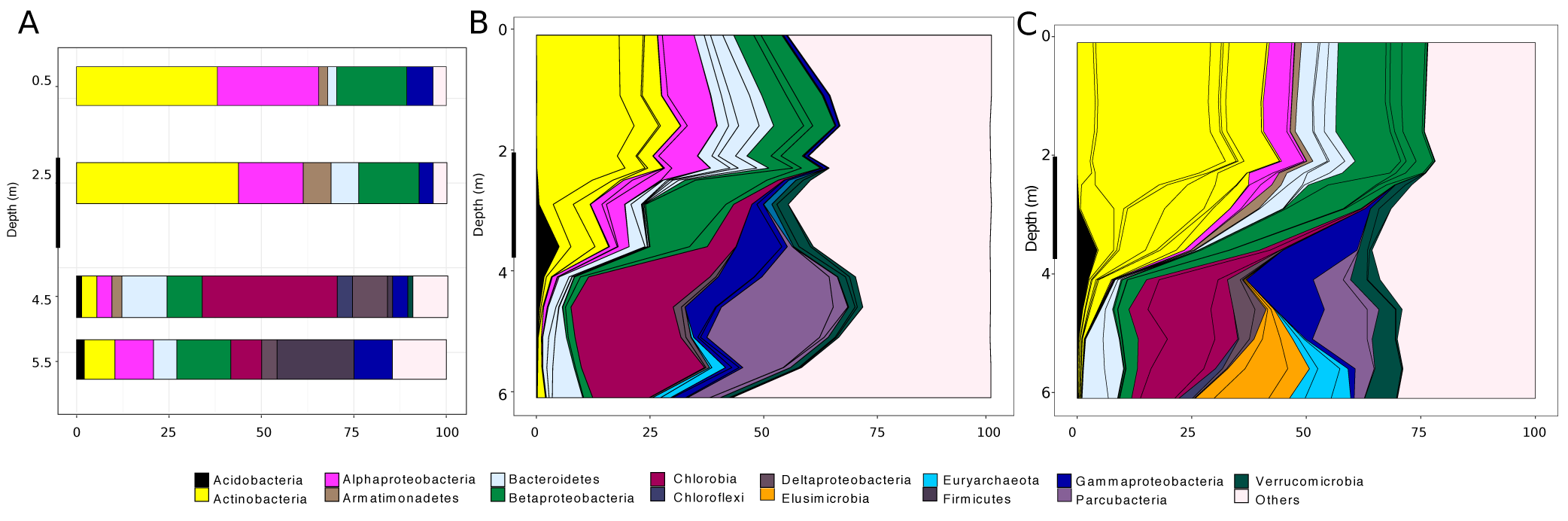
Microbial community composition in lake Alinen Mustajärvi based on A) 16S rRNA genes in 2008 (a clone library), B) 16S rRNA genes in 2013 (Illumina MiSeq), and C) metagenome assemble genomes from 2013 (Illumina HiSeq). In panels B and C different OTUs of the same phylum are separated by black lines. The black highlights on y-axis show the location of oxygen depletion zone.

The water column had a physico-chemical stratification with water temperature, oxygen (O_2_) and sulfate (SO_4_) concentrations decreasing with depth whereas the concentrations of CH_4_, DOC, ammonia (NH_4_), nitrite (NO_2_) and phosphate (PO_4_) were highest at the lake bottom (Fig. 3A-D). The microbial community was stratified into distinct layers with positive correlation between community composition in samples taken within 1 m distance of each other and negative correlations among samples taken with more than 2.6 m apart (Fig. 3E). While the bacterial community was homogenous throughout the epilimnion, below there was a sharp change coinciding with the decrease in oxygen concentration in the metalimnion and a subsequent succession of microbial taxa in the hypolimnion.

**Figure 3.**
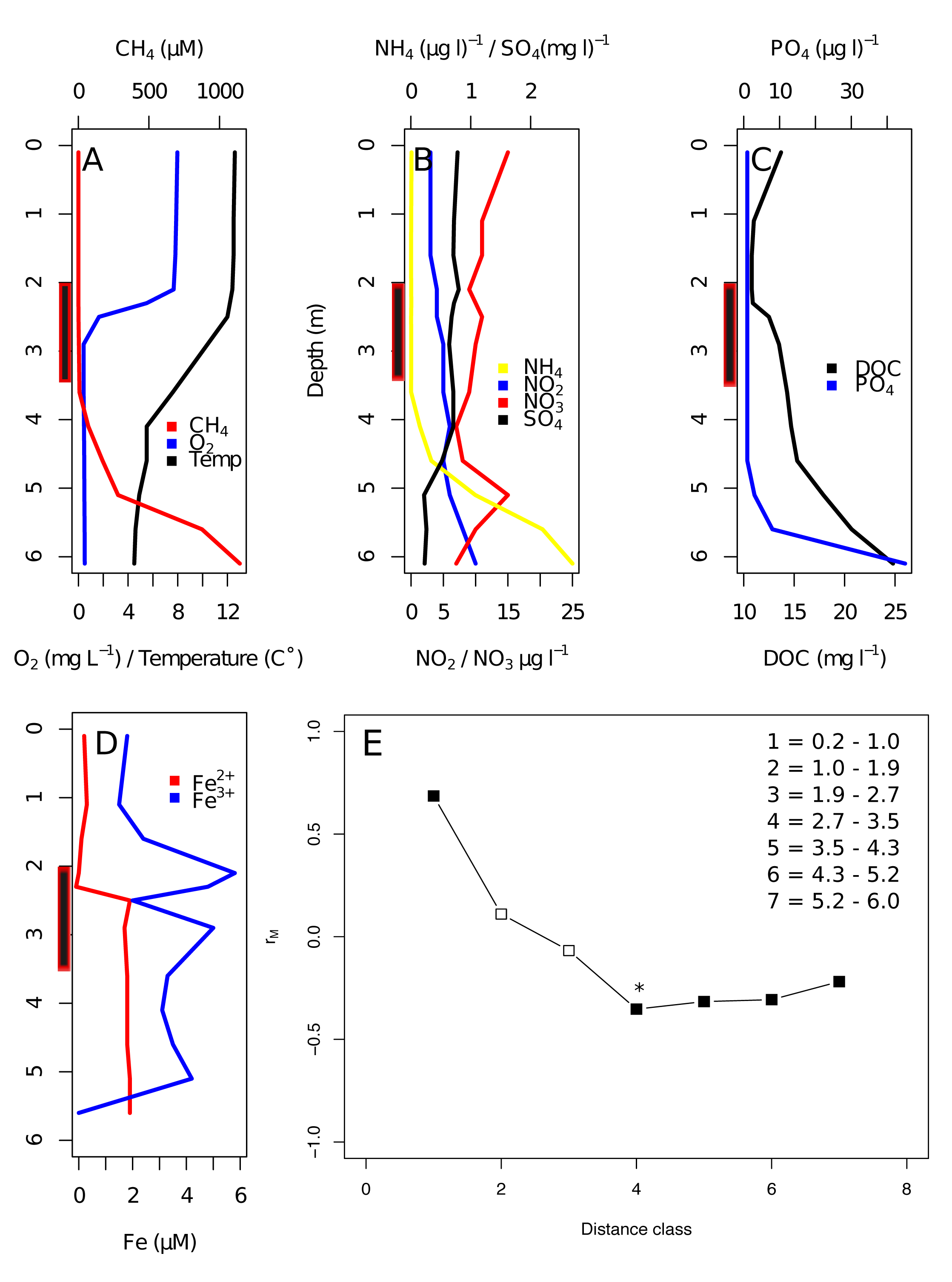
Environmental conditions in the lake in 2013. A) Concentration of O_2_ and CH_4_ and water temperature. Concentration of B) NH_4_, NO_2_, NO_3_ and SO_4_, C) PO_4_ and DOC, and D) Fe^2+^ and Fe^3+^. E) Bonferroni corrected Pearson correlations of the community composition (measured as Bray-Curtis distances) of the samples according to sampling distance. Black symbols designate significant correlations with p < 0.005 to all except distance 4, for which p = 0.006. The black highlights on the y-axis of panels A-D illustrate the oxygen depletion zone.

### Stratification of the microbial functional potential follows the redox gradient

The functional potential of the microbial community was estimated based on shotgun sequencing of the total DNA. Sequence coverage among samples (i.e. the proportion of reads from each of the samples that could be mapped to contigs) varied from 40.2 at the bottom to 75.5% in the surface, reflecting the diversity gradient in the lake (Table 1). Thus, our functional analysis may be missing some of the pathways present in the lake, but is expected to reflect the dominant metabolic potential in the water column. The abundances of different genes were normalized sample-wise using the abundance of 139 single copy genes (20). With this approach the abundance of the markers is expressed as Occurrences per Genome equivalent (GE), e.g. value of 1 suggests the presence of the gene in every genome of the sample. It should be noted that the values suggesting abundances of 1 and above could be explained by the presence of multiple gene copies in the genomes harboring these markers.

**Table 1.**
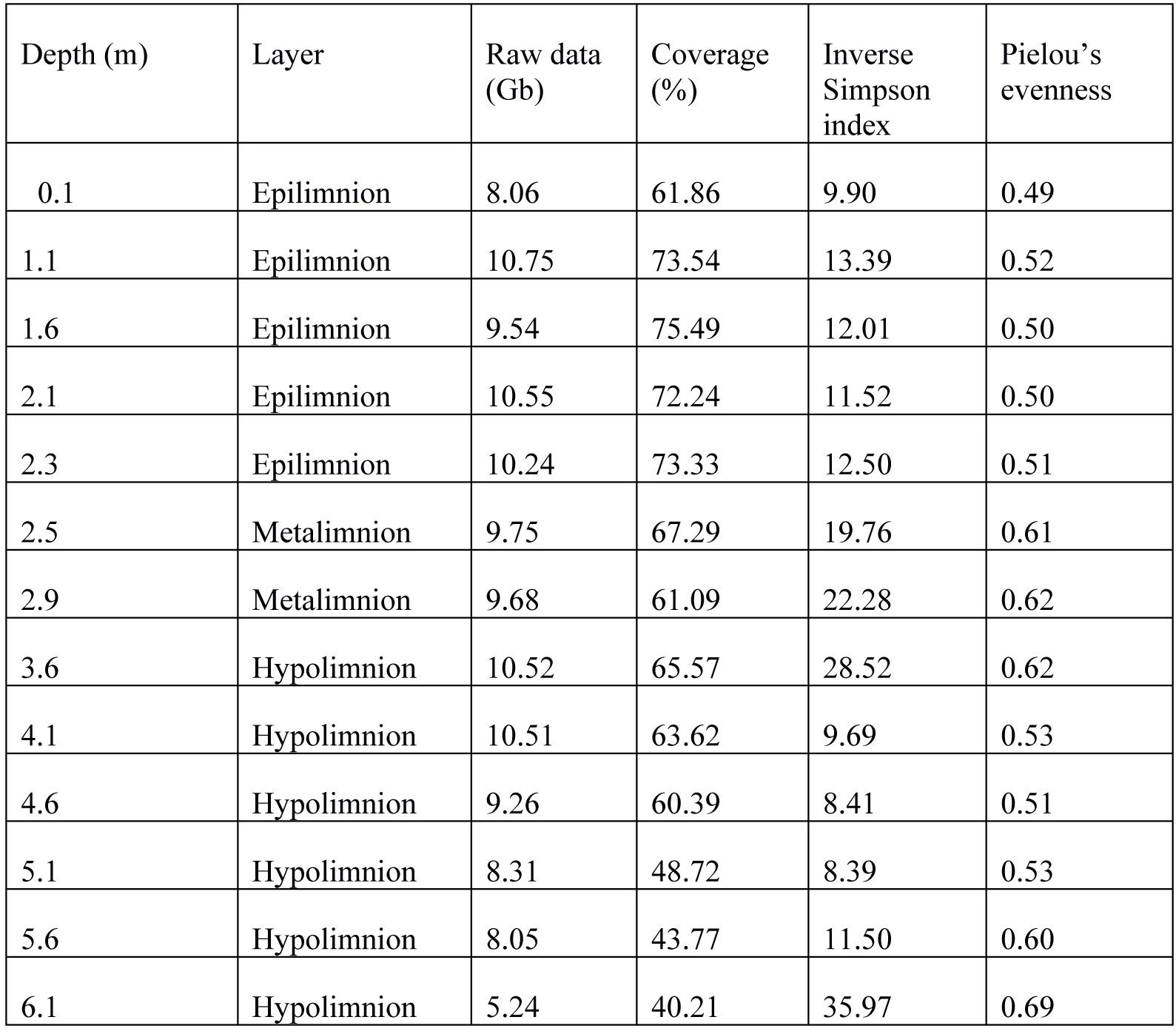
Sampling depth, water layer, data size, assembly coverage in each sample, and inverse Simpson index and Pielou’s evenness of the 16S rRNA OTU0.03 data.

| Depth (m) | Layer | Raw data (Gb) | Coverage (%) | Inverse Simpson index | Pielou's evenness |
| --- | --- | --- | --- | --- | --- |
| 0.1 | Epilimnion | 8.06 | 61.86 | 9.90 | 0.49 |
| 1.1 | Epilimnion | 10.75 | 73.54 | 13.39 | 0.52 |
| 1.6 | Epilimnion | 9.54 | 75.49 | 12.01 | 0.50 |
| 2.1 | Epilimnion | 10.55 | 72.24 | 11.52 | 0.50 |
| 2.3 | Epilimnion | 10.24 | 73.33 | 12.50 | 0.51 |
| 2.5 | Metalimnion | 9.75 | 67.29 | 19.76 | 0.61 |
| 2.9 | Metalimnion | 9.68 | 61.09 | 22.28 | 0.62 |
| 3.6 | Hypolimnion | 10.52 | 65.57 | 28.52 | 0.62 |
| 4.1 | Hypolimnion | 10.51 | 63.62 | 9.69 | 0.53 |
| 4.6 | Hypolimnion | 9.26 | 60.39 | 8.41 | 0.51 |
| 5.1 | Hypolimnion | 8.31 | 48.72 | 8.39 | 0.53 |
| 5.6 | Hypolimnion | 8.05 | 43.77 | 11.50 | 0.60 |
| 6.1 | Hypolimnion | 5.24 | 40.21 | 35.97 | 0.69 |

Despite the shallow light penetration depth (Fig. S2), markers indicative for phototrophy could be found throughout the water column (Fig. 4). The genes encoding for oxygen-evolving photosystem II, and aerobic anoxygenic photosynthesis (AAP) decreased rapidly with depth with combined abundance of these pathways at the surface being 1 GE. A second peak in photosynthetic potential was observed in the hypolimnion, where the abundance of the marker for anaerobic anoxygenic photosynthesis (bacterial photosynthesis) was up to 0.32 GE (Fig. S3).

**Figure 4.**
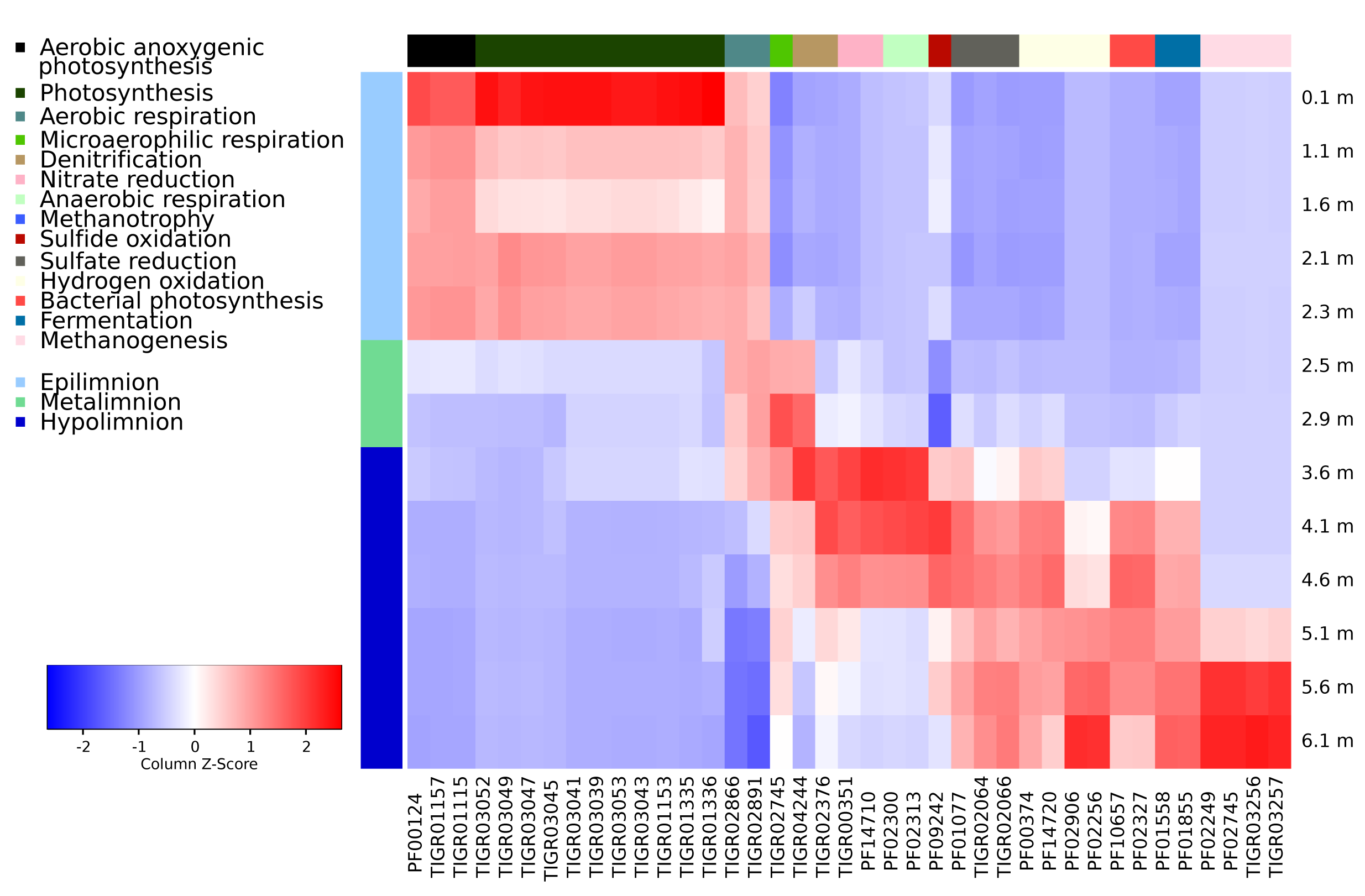
Heatmap visualizing the abundances of pfam/tigrfam markers related to energy metabolism (as given in z-score standardized per genome equivalent). Only those pfams/tigrfams that had significantly different abundance between different layers of the lake are displayed. Colors at the top of each of the columns reflect the function that the marker represents and colors on the left side of the heatmap illustrate different layers of the lake.

An iron gradient in the water column suggested that redox reactions related to iron could be important pathways for energy acquisition (Fig. 3D) and putative iron oxidizers (order *Ferrovales*) were abundant in the metalimnion. However, there were no specific markers for this pathway, thus while we have support for the potential for using iron as an energy source, we cannot visualize the trend for this pathway in the water column. For hydrogen oxidation the potential increased towards the lake bottom with the nickel dependent hydrogenase being more abundant right below the metalimnion and iron dependent at the lake bottom (Fig. 4), where the abundance of hydrogenases was close to 1 GE.

Based on their concentrations, the most important inorganic electron donors in the water column appeared to be CH4 in the upper hypolimnion, and sulfide/sulfur, that spanned almost throughout the hypolimnion. The peak of the marker indicative for sulfide/sulfur oxidation coincided with the maxima in phototrophic sulfur-oxidizing *Chlorobia*, a major community member in the hypolimnion. At this depth, the abundance of the markers for sulfide oxidation was 0.39 GE. The marker specific for sulfate reduction suggested highest potential in thelower hypolimnion. Markers for bacteria using anaerobic ammonia oxidation (ANAMMOX) were not detectable in the dataset. We could find markers for ammonia monooxygenase, but a manual inspection of the hits to these HMMs indicated that these were in fact to its paralog methane monooxygenase. Thus, these HMMs were used as markers for methanotrophy instead of nitrification.

The potential for aerobic respiration was stable from epilimnion to upper hypolimnion, whereas the highest potential for microaerophilic respiration was right above the depth where oxygen concentration dropped below detection limit and the potential for anaerobic respiration was highest in the upper hypolimnion. Potentials for using alternative electron acceptors for respiratory reactions followed the classic redox tower being nitrous oxide (N_2_O), NO_2_, NO_3_, Fe^3+^, SO_4_ and CO_2_ from metalimnion to the lake bottom. The profile suggested that the reactions in the denitrification pathway (reduction of nitrate to N_2_O or further to nitrogen gas N_2_) were divided between multiple organisms inhabiting different redox zones in the oxic-anoxic boundary layer, as the highest potential for different parts of the pathway were found at different depths (Fig. 4). Previous results on stratified lakes suggest that N_2_O commonly accumulates at the oxycline (21), as it is produced through both anaerobic nitrate reduction in the anoxic hypolimnion and aerobic nitrification in the oxic water layers. Furthermore, the N_2_O accumulation has been explained with the potential for nitrite reduction being higher than the potential for N_2_O reduction (9, 21). This is due to the expression of N_2_O reductase gene (nosZ) being more sensitive to oxygen than expression of nitrite reductase gene (nirK)(22). In our pfam/tigrfam profiles, the potential for nitrate reduction occurred in the upper hypolimnion and N_2_O reduction in the lower metalimnion, while markers for nitrification were sparse, suggesting that N_2_O dynamics is driven by anaerobic processes, potentially using NO_3_ as electron acceptor. The general patterns for Fe^3+^ and SO_4_ reduction could not be assessed due to lack of pathway specific HMMs, but these were investigated using metagenome assembled genomes (MAGs; Table S2).

### Unrecognized metabolic and chemoautotrophic potential in novel bacterial taxa

In general the microbial community had potential for three different pathways of autotrophic carbon fixation (Fig. 5). In the epilimnion, Calvin-Benson-Bassham cycle (CBB; also called reductive pentose phosphate pathway) was highly abundant, while reductive citric acid cycle (rTCA) and Wood-Ljungdahl pathway (WL; also called reductive acetyl CoA cycle) were mainly found in the hypolimnion. For the fourth carbon assimilation pathway present in the communities, 3- hydroxypropionate cycle (3HP), no specific HMMs could be found. However, protein annotations from Prokka suggested that multiple organisms possessed an almost full 3HP pathway, but certain genes in the pathway, such as mcr (malonyl CoA reductase), were not present in the dataset. The data suggested increasing autotrophic potential towards the bottom of the lake with a peak right below the oxycline and the highest potential at lake bottom with the total abundance of pathways related to carbon fixation close to 1 GE (Fig. 5).

**Figure 5.**
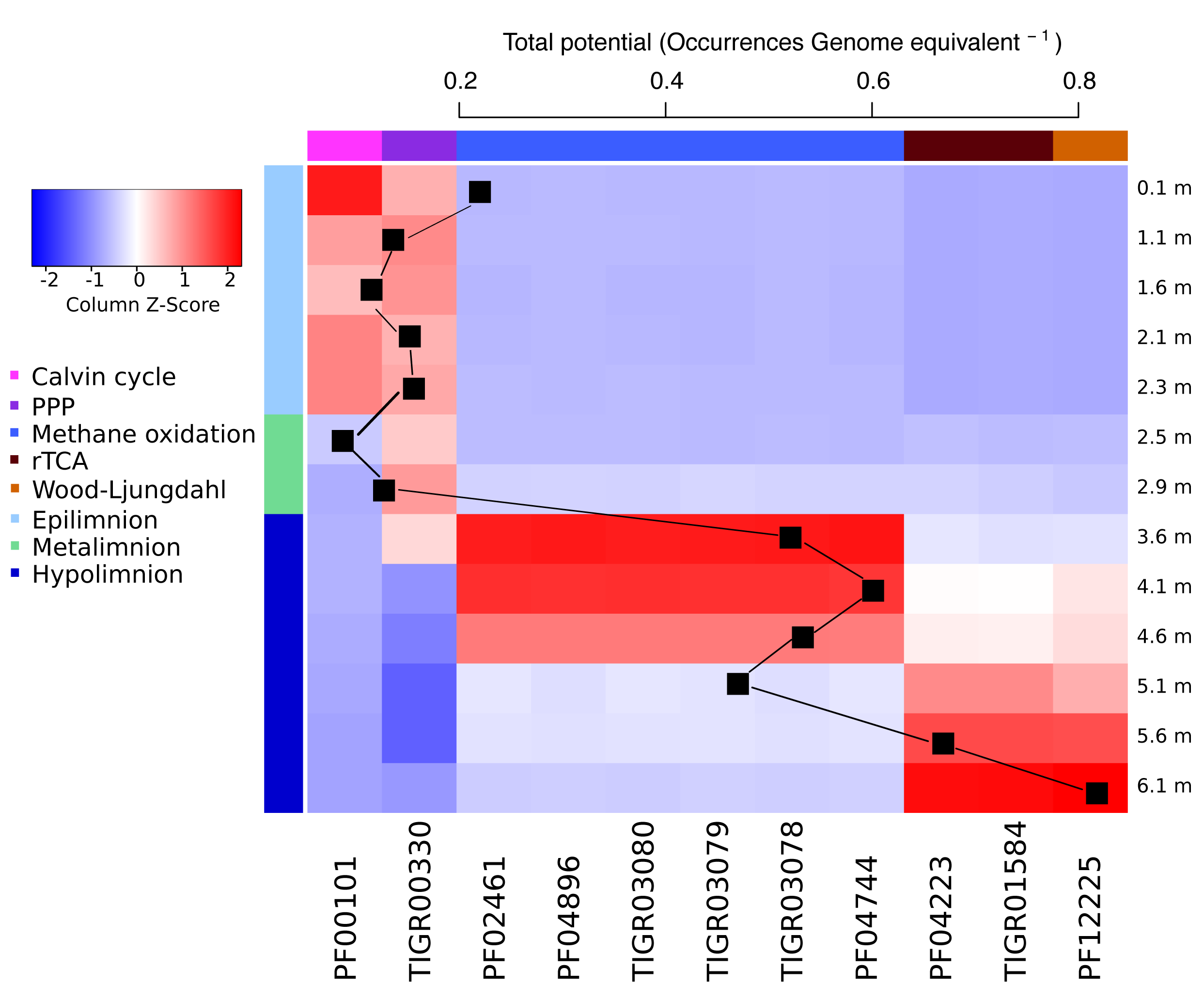
Heatmap visualizing the abundances of pfam/tigrfam markers (as given in z-score standardized per genome equivalent) related to carbon fixation and the total abundance of these genes in the dataset as a sum of the average abundances of the markers for each pathway, respectively. Only those pfams/tigrfams that had significantly different abundance between different layers of the lake are displayed. Colors at the top of each of the columns reflect the function that the marker represents and colors on the left side of the heatmap illustrate different layers of the lake.

We were able to construct a total of 270 metagenome assembled genomes (MAGs) of which 93 fulfilled our criteria for a high quality MAG (> 40 %completeness and < 4 % contamination). These were identified using PhyloPhlan (Fig. S4) (23). The high quality MAGs had the potential to use a wide range of different electron donors and acceptors and for a full range of microbial metabolic pathways from chemolithoautotrophy to photoheterotrophy (Table S2). Here we concentrate on the most abundant organisms with chemotrophic potential.

As stated above, the chemotrophic organisms appeared to be deriving their energy mainly from oxidation of sulfur, iron and hydrogen compounds. The most abundant MAGs with potential for sulfur oxidation (sox operon typically including genes soxACXYZ) included organisms related to *Chlorobia* (bin 4), *Polynucleobacter* (bins 15 and 120), *Comamonadaceae* (bin 108), *Ferrovales* (bin 139a), *Rhizobiales* (bin 139b) and *Acetobacteraceae* (bins 72 and 168). However, only the *Chlorobia* and *Ferrovales* MAGs had potential for autotrophy. *Chlorobia* was located in the lake hypolimnion and had a near-complete complex for oxidation of reduced sulfur compounds and also a near-complete CBB pathway for autotrophic carbon assimilation. Further, two other *Chlorobium* MAGs (bins 52 and 92) had dsr-operon, which is used in reverse in *Chlorobium* to oxidize sulfur (24). *Chlorobia* are known to be able to grow both autotrophically and mixotrophically (25). They derive the energy for autotrophy from the sunlight using sulfur oxidization to produce reducing equivalents for growth. Thus, they are photoautotrophic organisms rather than chemoautotrophs.

The other MAG with capacity for sulfur oxidation and inorganic carbon assimilation was a MAG closely related to betaproteobacterial *Ferrovales-order* (bin 139a). This MAG was 4.6 Mb in size (91.3% complete with 0.67% contamination) and included all of the genes in the CBB and rTCA pathways, suggesting the potential for autotrophy. While the MAG had potential for sulfur oxidation, it also encoded genes for iron oxidation: cytochrome cyc1, cytochrome-c oxidase (ctaCDE), ubiquinol-cytochrome-c reductase (petABC) and NADH:quinone oxidoraductase (nuoABCDEFGHIJKLMN). Further, the genome included an iron oxidase, which was located on the same contig as cytochrome cyc1, completing the pathway. This is consistent with the closest cultivated organism, iron oxidizing *Ferrovum mycofaciens*(26), which has previously been shown to thrive in acidic mine drainage (26). The MAG also included markers indicative of aerobic, anaerobic and microaerophilic respiration. Thus, the electron acceptor could be O_2_ when it is available. However, we could not identify any other electron acceptors for this organism. Thus, the genomic features together with the abundance distribution in the lake suggest that this organism is a chemoautotroph inhabiting suboxic to anoxic environments, using iron or sulfur oxidation as an energy source. In our data this organism was rather abundant in both 16S rRNA amplicon data, and in the metagenomes, but has not been previously found in boreal lakes. This particulate taxon was abundant in a narrow zone (between 2.5-3.6m), thus, a possible reason why it has been previously missed is the sparse sampling schemes of many experiments. Moreover, it represents a recently established bacterial order (26) and this taxon may have been classified as “uncultured *Betaproteobacteria”* in previous studies.

MAGs with autotrophic potential also included organisms that would appear to acquire energy by combining oxidation of hydrogen to sulfate or nitrate reduction. Hydrogenases found in the data represented FeFe - and NiFe-typehydrogenases as described in (27), with the latter type being more prevalent among the MAGs. For example, a MAG closely related to *Gallionellaceae* (bin 129) carried a 1e type hydrogenase, which is specifically used for electron input to sulfur respiration, and the MAG did have potential for sulfur reduction. It also had a full CBB pathway. Bin 129 was closely related to betaproteobacterial *Sideroxydans lithotropicus*, which has previously been found thrive in the same environment with *Ferrovales* (28). Similar to bin 129, this organism has the potential for CBB cycle, however, it has been suggested to derive its energy from iron rather than hydrogen oxidation (28). Another MAG, closely related to *Desulfobulbaceae* (bin 93), had a 1c type hydrogenase, also typically related to sulfate respiration, and an almost complete pathway for sulfate reduction. It also appeared to have the potential for CO2 fixation through reductive TCA cycle. The closest relative to this bin, *Desulfotalea psychrophila*, is also a sulfate reducer, but has been reported to be heterotrophic instead of autotrophic (29). However, *D. psychrophila* inhabits cold enviroenments, which is consistent with bin 93, which was most abundant in the deep layer of the lake where the water temperature is around 4 °C throughout the year.

In accordance with the redox potentials in the water column, and literature (30, 31), the reduction of nitrate to N2 was dispersed among multiple organisms. Candidatus Methyloumidiphilus alinensis (bin 10) (32) and a MAG closely affiliated with *Chrenotrix* (bin 149) both had a complete narGHIJ operon for nitrate reduction, and also genes for methane oxidation, as has been previously described for a member of *Methylobacter* family (33). Gene NosZ, coding for N_2_O reductase, was present in two high quality MAGs, which were taxonomically assigned to *Myxococcales* (bin 233) and *Bacteroidetes* (bin 64). However, these did not appear to be autotrophic organisms. NorCB operon, encoding NO reductase, was complete in two MAGs, in Candidatus M. alinensis and in a MAG affiliated to *Comamonadaceae* (bin 239). The latter was also carrying the potential for CBB cycle. The possible electron acceptors for the organism were sulfur and hydrogen. At the very bottom of the lake we could identify three archeal MAGs. Two of these were hydrogen oxidizing (hydrogenotrophic methanogenesis; bins 133 and 155, closely related to *Metanolinea* and *Methanoregula*, respectively), while the third one was using acetate (acetogenic methanogenesis; bin 74 closely related to *Methanosaeta*).

## Conclusions

Consistent with our expectations, the microbial community included a variety of different chemotrophic pathways. Further, the microbial community driving these processes contained abundant novel organisms with the potential for autotrophy. The assembly of the functional potentials was in accordance with redox potentials of the electron acceptors in the lake (Fig. 6). Further, consistent with our hypothesis, there was a diverse set of abundant chemoautotrophic organisms below the euphotic zone of the lake. The autotrophic community had an unexpected major community member closely related to *Ferrovales*, presumably thriving via iron oxidation. We also identified other novel autotrophs, such as an organism related to recently identified autotrophic *Sideroxydans lithotropicus*.

**Figure 6.**
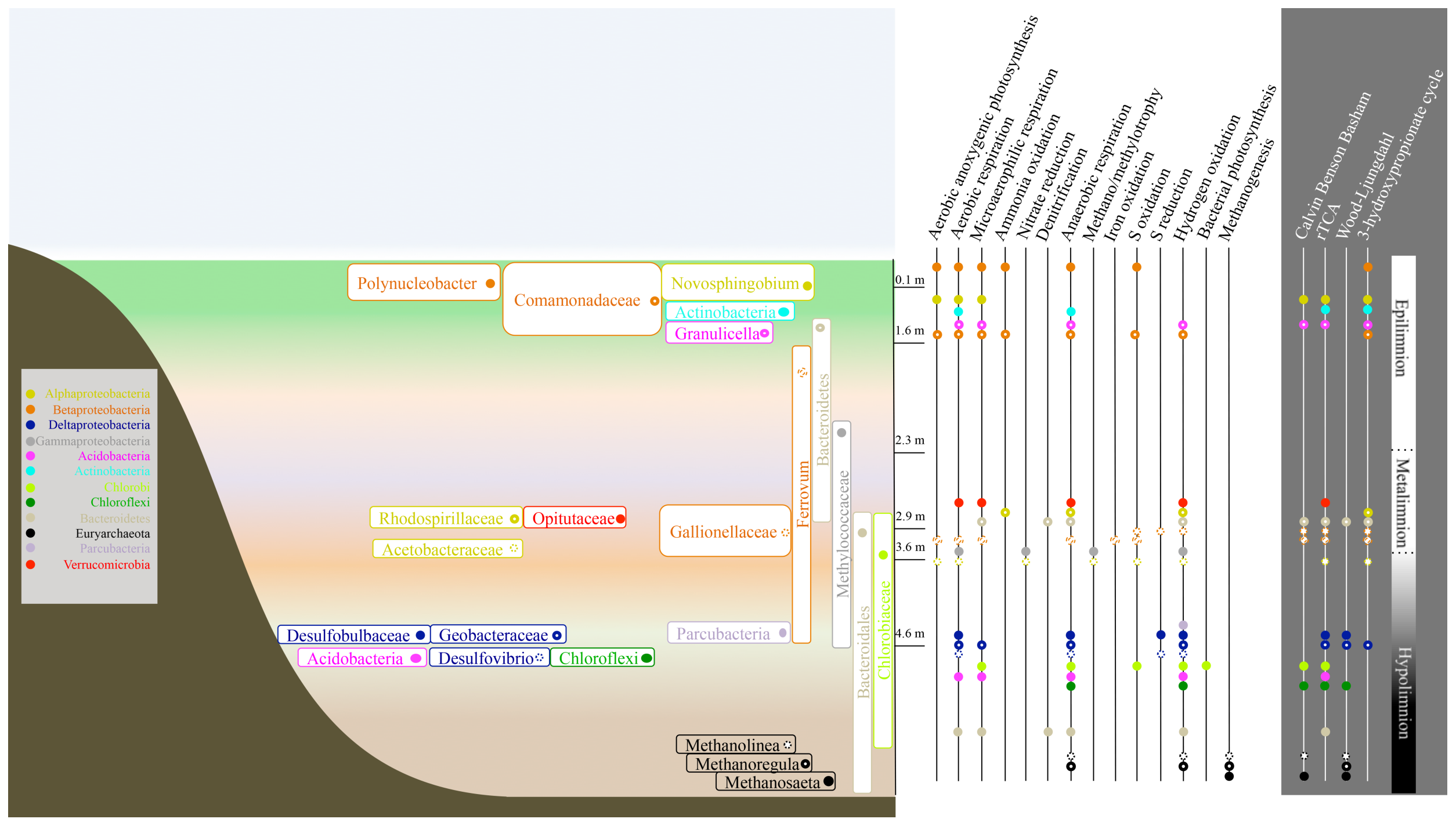
Redox reactions potentially driving the autotrophic processes in the lake and the most abundant organisms harboring these pathways. The colors represent the taxonomic annotation of the organisms at the phylum level or, in the case of Proteobacteria, the order. The height of the boxes visualizes the depth of themaximum abundance and in the case of multiple MAGs of the same taxon, the height of the box is covering the depths where the organisms were most abundant. Also, for the organisms with multiple MAGs with the same taxonomy, dominant pathways are displayed. Only the MAGs with marker genes specific for inorganic carbon fixation pathways are presented.

The fact that many of the autotrophic microbes were among the most abundant microbes in the lake and the abundance of photoautotrophic *Chlorobia*, emphasize the potential role of internal carbon cycling as a process that mitigates the flow of CO_2_ from boreal lakes to the atmosphere. Our results suggest that autotrophic iron, sulfur and hydrogen oxidizing microbes have a high potential to significantly contribute to inorganic carbon fixation in the lake. In fact, our measurements of inorganic carbon incorporation suggested that a significant amount of CO_2_ originating from degradation of autochthonous and terrestrial carbon can be re-incorporated into biomass in the poorly illuminated, anoxic layer of the lake. This is also well in line with results showing that chemolithoautotrophy significantly contributes to carbon and energy flow in meromictic lake Kivu (2), and with recent results regarding the autotrophic potential in boreal lakes (8). These processes are strongly dependent on the prevalent environmental conditions, which have been predicted to change following the warming of the climate. Thus, our results suggest that if the predicted alterations in lake environment, such as changes in mixing patterns, should happen, we may expect reorganization of the metabolic processes in the lake, which would have unknown implications to the carbon flow in the water column.

## Materials and Methods

### Site description and sampling

The study lake, Alinen Mustajärvi, is situated in southern Finland (61°12’N, 25°06’E). It is a 0.007 km^2^ head-water lake with maximum depth 6.5 m and an estimated volume of 31 × 10^3^ m^3^. The catchment area is <0.5 km^2^ and it consists of>90%coniferous forest and <10% peatland. The lake is characterized by steep oxygen stratification during summer and also during ice cover period, which lasts from late November until late April. The stratification is disrupted by regular autumn and irregular spring mixings. As such Alinen Mustajärvi is a representative for the millions of lakes and ponds in the arctic and boreal zones.

The metagenome sampling was conducted in the beginning of September 2013, at the end of stratification period. The lake was sampled at 13 depths; the oxic epilimnion was sampled at 0.1, 1.1, 1.6, 2.1 and 2.3 m; the metalimnion at 2.5 and 2.9 and the hypolimnion at 3.6 m, 4.1, 4.6, 5.1, 5.6 and 6.1 m. Water samples were taken with a 20-cm-long acrylic tube sampler (Limnos vol. 1.1 L) and subsequently analyzed for nutrients (NO_2_/NO_3_, PO_4_, NH_4_, total N, total P, SO_4_), gasses (CH_4_, CO_2_) and dissolved organic carbon (DOC) concentration. Nutrient analyses were conducted using standard methods (http://www.sfs.fi/). Gas and DOC analyses were done as in (34) and iron as in (35). The dark carbon fixation was measured in 2008 as an increase or decrease of dissolved inorganic carbon (DIC) concentration during 24 h incubation at the depth of 3 m in two foil-covered 50-ml glass stoppered BOD bottles. The measurements were conducted every second week from beginning of May until end of October and the changes in DIC concentration were analyzed according to (36). In August 2008 a clone library was created from depths 0.5, 2.5, 4.5 and 5.5 m, consisting of a total of 303 sequences (37).

The samples for metagenomic analysis of lake microbiota were taken by filtering water through a 0.2 μm polycarbonate filters which were then frozen at −78 °C until further analysis. The DNA was extracted from the filters using Mobio PowerSoil DNA extraction kit (MO BIO Laboratories). Sample preparation for 16S rRNA gene analysis and the following sequence processing were conducted as previously described (38).

Shotgun metagenomic libraries were prepared from 10 ng of genomic DNA. First, the genomic DNA was sheared using a focused-ultrasonicator (Covaris E220) and subsequently, sequencing libraries were prepared with the Thruplex FD Prep kit from Rubicon Genomics according to the manufactures protocol (R40048-08, QAM-094-002). Library size selection was made with AMPure XP beads (Beckman Coulter) in 1:1 ratio. The prepared sample libraries were quantified using KAPA Biosystem’s next-generation sequencing library qPCR kit and run on a StepOnePlus (Life Technologies) real-time PCR instrument. The quantified libraries were then prepared for sequencing on the Illumina HiSeq sequencing platform with a TruSeq paired-end cluster kit, v3, and Illumina’s cBot instrument to generate a clustered flowcell for sequencing. Sequencing of the flow cell was performed on the Illumina HiSeq2500 sequencer using Illumina TruSeq SBS sequencing kits, v3, following a 2x100 indexed high-output run protocol.

The sequencing produced a total of 120.5 Gb of sequence data. The raw data has been deposited to NCBI Sequence Read Archive under accession number SRP076290. Reads were filtered based on their quality scores using sickle (version v1.33) (39) and subsequently assembled with Ray (version v2.3.1) (40). Assembled contigs from kmer sizes of 51, 61, 71 and 81 were cut into 1000 bp pieces and scaffolded with Newbler (454 Life Sciences, Roche Diagnostics). Mapping of the original reads to the Newbler assembly was done using bowtie2 (version v2.15.0) (41),while duplicates were removed using picard-tools (version 1.101; https://github.com/broadinstitute/picard), and for computing coverage, bedtools (42) was used. Details on the assembly results are presented in Table 1. The data was then normalized using the counts of 139 single copy genes as previously described (20). Assembled contigs were binned with MetaBAT (version v0.26.3) (43) to reconstruct genomes of the most abundant lake microbes (metagenome assembled genomes (MAGs)). The quality of the MAGs was evaluated using CheckM (version v1.0.6) (44). Cutoffs for high quality MAGs were set to ≥ 40% for completeness and ≤ 4% for contamination. The placement of the MAGs in the microbial tree of life was estimated using Phylophlan (version v1.1.0) (23).

The functional potential of the metagenomes was assessed from assembled data using the hidden Markov models (HMM) of the Pfam and TIGRFAM databases (45, 46) and the HMMER3 software (version v3.1b2) (47). Special attention was paid to pathways linked to energy metabolism and carbon cycle. To assure pathway specificity, marker HMMs were chosen to be unique to specific pathways (Table S1). Normalized coverage information of the contigs combined with HMMs of specific marker genes (Table S1) was used to predict protein domains related to energy metabolism and carbon incorporation to biomass. Only marker genes that were found to exhibit a significantly different distribution between layers are reported (p-values in Table S1). All of the MAGs were also annotated using Prokka (version v1.11) (48). The metabolic potentials of all high quality MAGs were evaluated based on Prokka annotations (Table S2). In case of novel functional combinations, such as combination of AAP to CBB, the contigs were blasted against the NCBI nr database to verify that the closest relatives of the overall genes in these contigs were matching to the Phylophlan annotation. MAGs with special functional properties were also visualized with non-metric multidimentional scaling to check the placement of the contigs with markers within all the contigs comprising the MAG in question. All statistical analyses were done using R software (http://www.R-project.org(49)) and packages vegan (50) and mpmcorrelogram (51). Differences in the abundance of marker HMMs between layers was tested using permutation test (1000 permutations) on the t-statistics with package MetagenomeSeq (52).

## Acknowledgements

We thank Sainur Samad for his help with processing the DNA samples. Lammi biological station is acknowledged for facilities and equipment during the sampling as well as for nutrient analyses. Sequencing was performed at the SNP & SEQ technology platform at Science for Life Laboratory, Uppsala. IT Center for Science (CSC; Espoo, Finland) and Uppsala Multidisciplinary Center for Advanced Computational Science (UPPMAX; Uppsala, Sweden) provided the computational and data storage resources. Funding was provided by the Academy of Finland (Grant Number 265902 to SP, and for 2008 sampling grant number 114604 to Roger I Jones), the Tryggers Foundation (Grant to SP and AE), the Swedish Research Council VR (Grant 2012-4592 to AE) and the Swedish Foundation for strategic research (Grant ICA10-0015 to AE).

The authors declare no competing financial interests.

## Supplementary Tables

**Supplementary Table S1.** A table listing HMMs used to compare the abundance of different energy and carbon assimilation pathways between different depths layers and p-values of these comparisons.

**Supplementary Table S2.** Characteristics of the high-quality MAGs. Bin Id, taxonomic annotation by i) manual annotation from a phylogenetic tree from PhyloPhlan and ii) PhyloPhlan automatic annotation, size of the MAG, average proportion of the MAG among all samples, depth of the maximum abundance and the proportion in this depth, completeness, contamination and strain heterogeneity of the MAG and number of marker genes for different energy and carbon assimilation pathways in the MAG. Presence of key genes of each of the pathway is shown with letters. AAP: M = pufM; methanogenesis: M = mcrA; CBB: R = RuBisCo; PPP: Z = glucose-6-phosphate dehydrogenase, P = 6-phosphogluconolactonase, G = 6-phosphogluconate dehydrogenase; rTCA: C = citrate lyase, O = oxoglutarate synthase, P = pyruvate synthase; WL: C = CO dehydrogenase; 3HP: A = acetyl-CoA carboxylase, P = propionyl-CoA carboxylase. For hydrogenases sub groups are presented as in D. Søndergaard, C.N. Pedersen, and C Greening. Sci Rep 6:34212,2016.

## Supplementary Figures

**Supplementary Figure S1.** Change in dissolved inorganic carbon concentration during 24 h dark incubations at 3 m depth in 2008-2010. In 2008 measurements were done biweekly, in 2009 monthly and in 2010 three times during the open water season.

**Supplementary Figure S2.** Photosynthetically active radiation in the lake water column.

**Supplementary Figure S3.** The average abundance of marker genes for different pathways related to energy generation.

**Supplementary Figure S4.** Phylogenetic tree of the Metagenome Assembled Genomes depicting their taxonomic placement in the tree of life.

